# Host membrane cholesterol constrains constitutive signaling of the oncogenic KSHV GPCR ORF74

**DOI:** 10.64898/2026.09.22.753489

**Authors:** Jun Bae Park, Amita Rani Sahoo, Annabelle Ryu, Matthias Buck, Jae U. Jung

**Author notes:** These authors contributed equally (Jun Bae Park, Amita Rani Sahoo). Correspondence (Jun Bae Park) and (Jae U. Jung).

## Abstract

Cholesterol is a major structural component of the plasma membrane and a key allosteric regulator of G protein-coupled receptors (GPCRs), yet its role in controlling constitutive receptor activity remains poorly understood. Virally encoded GPCRs provide an ideal system to address this question because many exhibit constitutive signaling that promotes viral persistence and pathogenesis, although how excessive receptor activation is restrained remains unknown. Here, we identify a previously unrecognized cholesterol-dependent allosteric mechanism by which the oncogenic Kaposi’s sarcoma-associated herpesvirus (KSHV) GPCR ORF74 constrains its constitutive activity. CryoEM structural analysis reveals a cholesterol-binding pocket formed by transmembrane helices 3, 5, and 6 that is present only in the inactive receptor. Cholesterol binding restrains the outward movement of transmembrane helix 6, stabilizes the inactive conformation, and suppresses spontaneous receptor activation, as supported by molecular dynamics simulations. Mechanistically, replacement of the canonical DRY motif with a non-canonical VRY motif exposes the conserved R143^3.50^ residue, creating a membrane-facing cholesterol-binding interface that couples membrane cholesterol to receptor conformational control. Consistent with this model, cellular cholesterol depletion enhances ORF74 signaling, whereas disruption of cholesterol binding impairs receptor stabilization. Together, our findings uncover a previously unrecognized mechanism by which a viral GPCR exploits host membrane cholesterol to regulate its constitutive activity, suggesting that persistent viruses optimize constitutive signaling by coupling receptor activity to host lipid-dependent allosteric regulation.

**SIGNIFICANCE:** Constitutively active viral G protein-coupled receptors (GPCRs) promote viral persistence and oncogenesis, yet how excessive signaling is restrained has remained unknown. We identify a cholesterol-binding allosteric pocket that stabilizes the inactive conformation of the Kaposi’s sarcoma-associated herpesvirus GPCR ORF74, thereby suppressing constitutive receptor activity. Unexpectedly, the non-canonical VRY motif previously linked to constitutive signaling also creates the cholesterol-responsive interface, coupling intrinsic receptor activity to the host membrane environment. These findings reveal a mechanism by which a persistent virus exploits host membrane cholesterol to optimize rather than maximize signaling output and establish membrane cholesterol as a direct regulator of constitutive GPCR signaling with potential therapeutic implications.

## INTRODUCTION

G protein-coupled receptors (GPCRs) constitute the largest family of cell-surface signaling receptors and regulate virtually every aspect of animal physiology(1). Although GPCR activation has traditionally been viewed as a process governed by extracellular ligands and intracellular signaling proteins, it is increasingly recognized that the surrounding membrane actively participates in receptor regulation(2). Among membrane lipids, cholesterol has emerged as a major allosteric regulator capable of reshaping GPCR conformational landscapes, stabilizing inactive or active receptor states, altering signaling efficacy, influencing ligand binding, and organizing receptors within specialized membrane microdomains(2–5). Despite growing appreciation of cholesterol as a regulator of GPCR function, how membrane cholesterol controls constitutive receptor activity and whether this mechanism has been exploited during receptor evolution remains poorly understood.

Virally encoded GPCRs provide a unique opportunity to address this question. Many large DNA viruses have acquired host GPCR genes during evolution and subsequently remodeled them to promote viral persistence, immune evasion, and pathogenesis(6–8). Unlike their cellular counterparts, viral GPCRs frequently exhibit constitutive signaling, allowing continuous activation of downstream pathways independent of ligand binding(6–8). Constitutive signaling provides clear advantages by promoting cell survival, proliferation, inflammation, and viral persistence(9–12). However, persistent viruses are unlikely to benefit from unrestricted receptor activation because excessive signaling could compromise viral fitness by disrupting cellular homeostasis, enhancing immune surveillance, or reducing the long-term survival of infected cells(13, 14). How viral GPCRs maintain this balance between persistent signaling and controlled receptor activity remains an unresolved question.

One of the well-characterized viral GPCRs is ORF74 encoded by Kaposi’s sarcoma-associated herpesvirus (KSHV/HHV-8), the etiologic agent of Kaposi’s sarcoma, primary effusion lymphoma, and multicentric Castleman disease(15–17). ORF74 is a class A GPCR that displays exceptionally high constitutive activity, while retaining responsiveness to multiple chemokine ligands and diverse G proteins(6, 9, 18–21). Persistent activation of angiogenic, inflammatory, and proliferative signaling pathways by ORF74 is sufficient to drive endothelial transformation and Kaposi’s sarcoma-like lesions in vivo(20, 22) establishing ORF74 as one of the principal viral oncogenes and an attractive therapeutic target. Yet how ORF74 maintains robust constitutive signaling while avoiding excessive receptor activation remains incompletely understood.

In our recent work, we determined cryoEM structures of ORF74 in inactive apo and CXCL1/Gi_trimer_-bound active conformations, revealing how the receptor accommodates diverse chemokines while maintaining unusually high basal signaling activity(23). Structural and computational analyses demonstrated that ORF74 spontaneously samples active-like conformations, providing a molecular explanation for its constitutive signaling. This analysis further identified replacement of the highly conserved DRY motif of class A GPCR with a non-canonical VRY motif as a key determinant of the exceptional constitutive activity of ORF74(23). Despite these observations, however, the apo receptor unexpectedly adopted a canonical inactive conformation despite its strong intrinsic propensity for activation. This observation revealed a fundamental paradox: how can one of the most constitutively active GPCRs remain structurally stabilized in an inactive state?

A clue emerged from the inactive ORF74 structure, which contained an unassigned density embedded within the transmembrane region. More broadly, cholesterol has been shown to stabilize distinct conformational states in several cellular GPCRs(24–26), yet whether host membrane cholesterol similarly regulates virally encoded GPCRs remains largely unexplored. Because viral GPCRs have evolved signaling properties distinct from those of cellular receptors, host membrane cholesterol may provide an additional layer of allosteric regulation that constrains receptor activity. This possibility is particularly intriguing for ORF74, whose exceptionally high constitutive activity would be expected to require mechanisms that prevent excessive activation.

Here, we identify a previously unrecognized cholesterol-binding allosteric pocket that stabilizes the inactive conformation of ORF74 by restraining the outward movement of transmembrane helix 6, thereby limiting constitutive receptor activation. Unexpectedly, cholesterol binding is enabled by a non-canonical VRY motif that exposes the conserved R143^3.50^ residue, coupling membrane cholesterol to receptor conformational control. These findings suggest that the structural adaptation underlying constitutive signaling simultaneously creates a host cholesterol-dependent allosteric mechanism that constrains excessive receptor activation. More broadly, our work reveals how viral GPCRs can exploit conserved host membrane components to optimize signaling output, providing a conceptual framework for host lipid-dependent regulation of constitutively active receptors.

## RESULTS

### A membrane-facing cholesterol-binding pocket stabilizes the inactive ORF74 conformation

During re-examination of our previously determined inactive ORF74 cryoEM structure (EMD-43717, 2.89 Å resolution)(23), we identified a well-defined non-protein density embedded within the transmembrane region that had not been assigned in the original model. To accurately define this feature, we rebuilt and refined the structure, yielding a high-quality model with excellent agreement between the atomic coordinates and cryoEM density (overall Q-score = 0.61, ***SI Appendix* Fig. 1A**). The transmembrane helices exhibited particularly high local Q-scores, consistent with the excellent map quality in this region. As reported previously(23), the structure adopts an antiparallel dimeric arrangement, and the two protomers are nearly identical (Cα-backbone RMSD = 0.29 Å) (***SI Appendix* Fig. 1B**).

The unassigned density was consistently observed at an equivalent position in both ORF74 protomers, suggesting the presence of a specifically bound sterol molecule. Based on its characteristic size and shape, cholesteryl hemisuccinate (CHS), a cholesterol analog present throughout protein purification and cryoEM sample preparation, was modeled into the density. The fitted CHS molecule showed excellent agreement with the cryoEM map (Q-score = 0.58), supporting assignment of the feature as a sterol-binding site (***SI Appendix* Fig. 1A and B**). This assignment resolves the previously unexplained density in the inactive ORF74 structure and identifies a membrane-facing lipid-binding site that may contribute to stabilization of the inactive receptor.

The CHS molecule occupies a pocket located on the cytoplasmic side of the transmembrane domain (***SI Appendix*. Fig. 1C**). The pocket is formed by TM3, TM5, and TM6, creating a concave hydrophobic cavity that accommodates the sterol ring, whereas the hemisuccinate moiety extends toward intracellular loop 3 (ICL3), where it forms electrostatic interactions with positively charged residues. The observed binding pose closely resembles physiologically relevant cholesterol-binding modes reported in other membrane proteins(8, 27), supporting CHS as a structural surrogate for native cholesterol.

The location of this sterol-binding pocket immediately suggested a mechanism for receptor regulation because it directly contacts TM6, the transmembrane helix that undergoes the largest conformational rearrangement during activation of class A GPCRs(28). To determine whether cholesterol binding is coupled to receptor activation, we compared the inactive ORF74 structure with the previously reported active-state structure (PDB: 9EJC)(23). Whereas the TM3-TM5-TM6 pocket is well defined in the inactive receptor, outward displacement of TM6 in the active conformation collapses the cavity and eliminates the sterol-binding site (**Figure 1A** and **B**). Thus, cholesterol binding is structurally compatible only with the inactive receptor conformation, suggesting that cholesterol stabilizes ORF74 by restricting TM6 movement.

**Figure 1.**
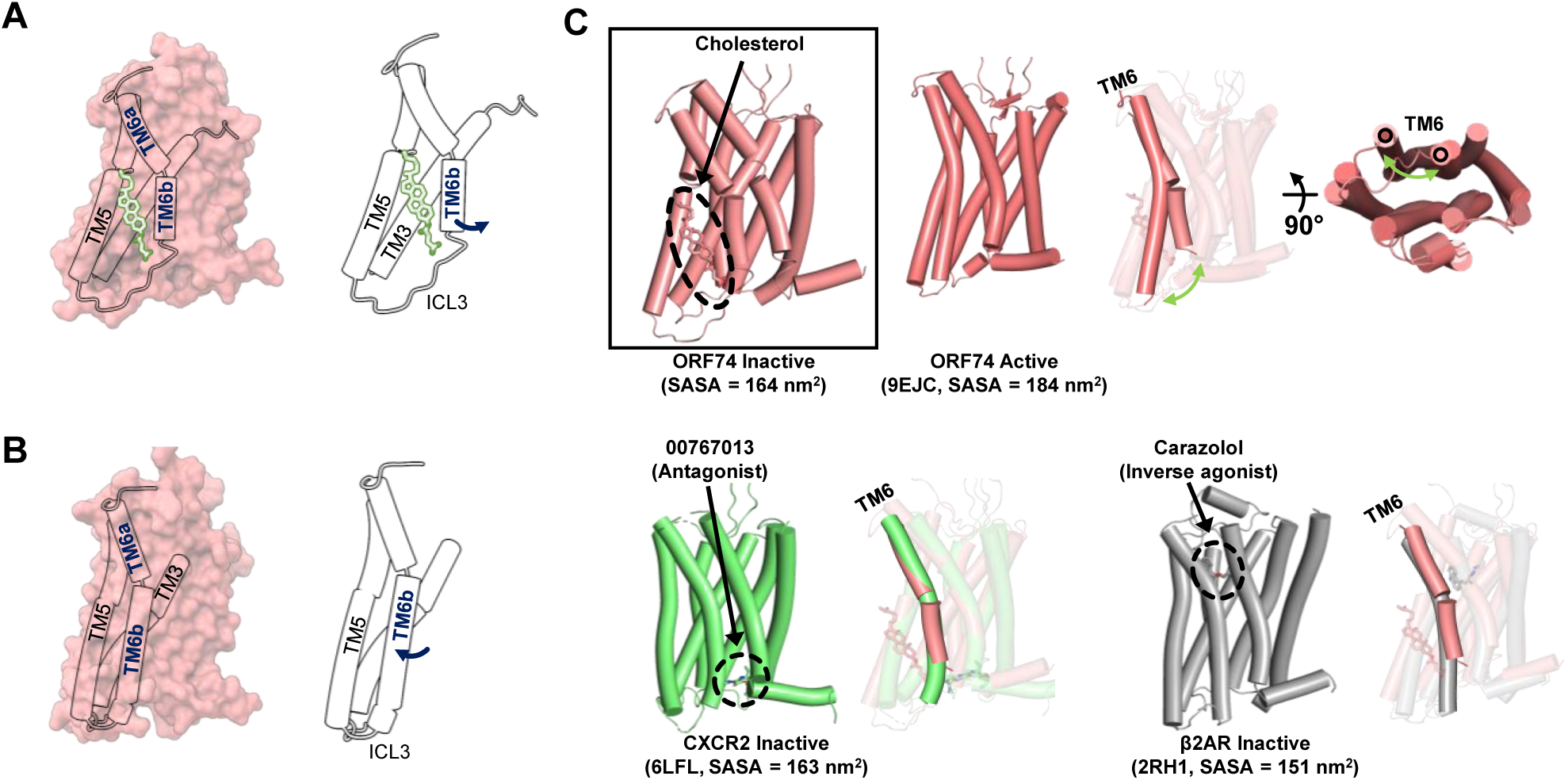
Cholesterol-binding pocket is uniquely formed in the inactive conformation of ORF74. **(A)** CryoEM structure of inactive ORF74 showing cholesterol (modeled as CHS) bound within a hydrophobic pocket formed by transmembrane helices (TM) 3, 5, and 6. Surface representation (left) and corresponding cartoon view (right) highlight the location of the cholesterol-binding pocket and the movement of TM6 that permits pocket formation. **(B)** Side view of the active ORF74 structure highlighting the spatial relationship between the cholesterol-binding pocket and TM6. Surface (left) and cartoon (right) representations show that movement of TM6 seals the cholesterol-binding pocket, restricting cholesterol access. **(C)** Comparison of TM6 conformations in class A GPCRs. Cholesterol binding in ORF74 is restricted to the inactive conformation, in which TM6 adopts a position similar to that of inactive CXCR2 and β2AR, supporting cholesterol-mediated stabilization of the inactive state. Pocket SASA is indicated for each structure.

To determine whether this sterol-binding mode is conserved among class A GPCRs, we compared ORF74 with representative cholesterol-bound GPCR structures compiled in a recent comprehensive survey(29). Among the receptors examined, the closest sterol-binding position was observed in CX3CR1(30) (***SI Appendix* Fig. 2A**). However, while cholesterol in CX3CR1 primarily interacts with TM5, the ORF74 pocket simultaneously engages TM3, TM5, and TM6. Direct participation of TM6 therefore distinguishes the ORF74 pocket from previously characterized cholesterol-binding sites and uniquely positions cholesterol to regulate the principal conformational switch governing GPCR activation.

To further define the conformational state associated with cholesterol binding, we compared ORF74 with representative inactive-state structures of its sequence and functional homolog CXCR2 and the prototypical class A GPCR β2-adrenergic receptor (β2AR)(31–33). The TM6 conformation of cholesterol-bound ORF74 closely resembles that observed in antagonist-bound CXCR2 and inverse agonist-bound β2AR, both of which represent canonical inactive-state structures (**Figure 1C**). Collectively, these findings identify a previously unrecognized TM3-TM5-TM6 cholesterol-binding pocket that is unique among currently characterized GPCR cholesterol-binding sites and suggest that membrane cholesterol stabilizes ORF74 by maintaining the receptor in a canonical inactive conformation.

### Cholesterol stabilizes the inactive conformation of ORF74 by restraining TM6 movement

The structural analyses described above suggested that cholesterol binding may stabilize the inactive conformation of ORF74 by restricting movement of TM6, the transmembrane helix that undergoes the largest conformational rearrangement during activation of class A GPCRs(31). Outward movement of TM6 opens the intracellular pocket for G protein and β-arrestin binding. To investigate how membrane cholesterol influences the conformational dynamics of ORF74, we first performed molecular dynamics simulations of the inactive ORF74 structure embedded in a membrane containing 30% cholesterol, which closely approximates the cholesterol content of the human plasma membrane(34). These simulations were compared with our previously reported trajectories of the inactive and active ORF74 structures in cholesterol-free membranes(23). Representative structures after 200 ns revealed markedly different conformational behaviors depending on membrane cholesterol content (**Figure 2A**). In the absence of cholesterol, TM6 underwent a gradual outward displacement toward an active-like conformation, accompanied by deformation of the cholesterol-binding pocket. This behavior is consistent with the intrinsic constitutive activity of ORF74 and its tendency to spontaneously sample active-like conformations(23). In contrast, the inactive receptor remained structurally stable in the 30% cholesterol membrane, TM6 maintained its inactive orientation and exhibited only a slight inward movement throughout the simulation. These observations suggest that membrane cholesterol stabilizes the inactive receptor by limiting the outward movement of TM6, the hallmark structural transition associated with GPCR activation.

**Figure 2.**
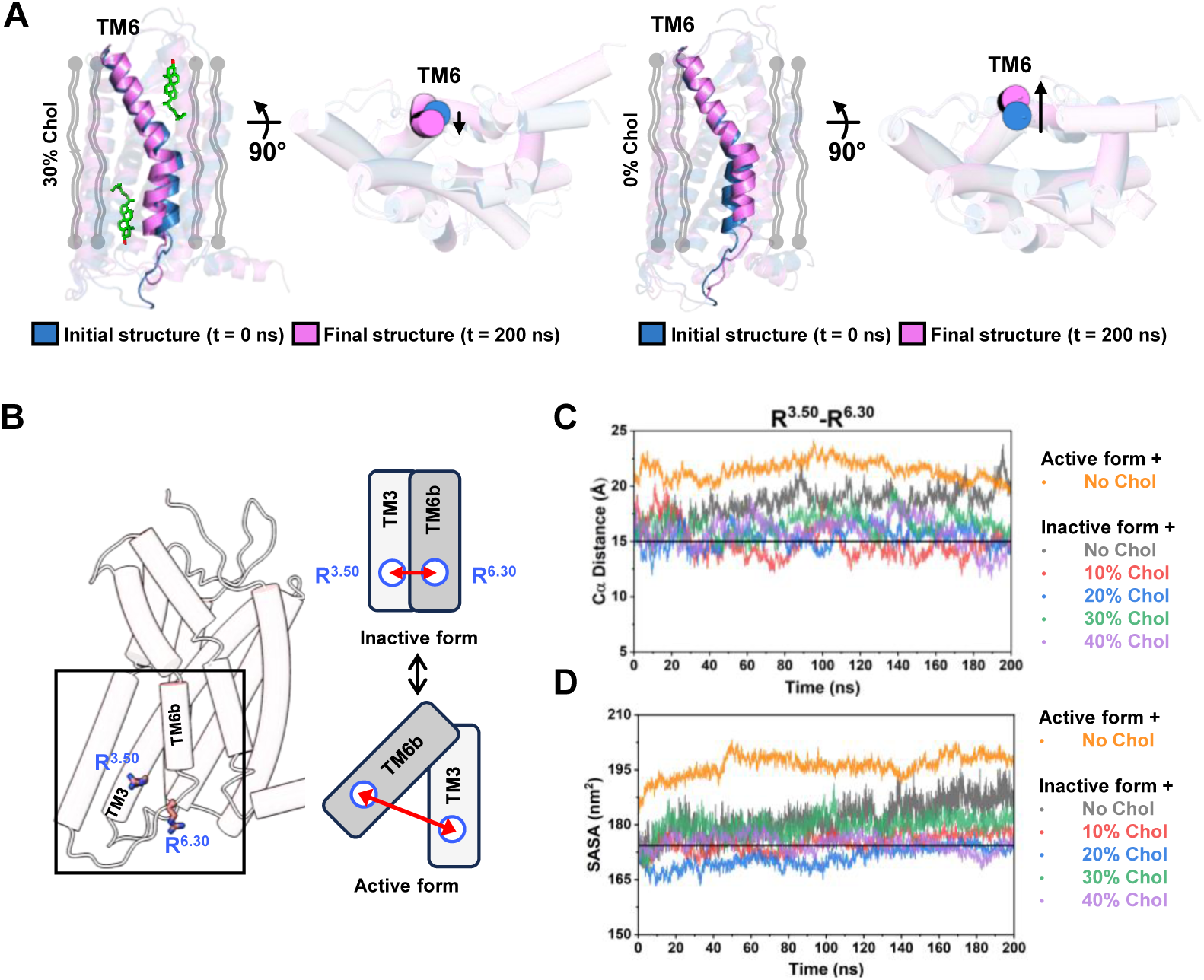
Cholesterol stabilizes the inactive conformation of ORF74 during molecular dynamics simulations. **(A)** Representative structures of the inactive ORF74 receptor after 200 ns molecular dynamics simulations in membranes containing 30% cholesterol (left) or lacking cholesterol (right). The initial structure (light blue, t = 0 ns) is superimposed with the final structure (purple, t = 200 ns). Side and intracellular views show that cholesterol stabilizes the inactive position of TM6, whereas cholesterol depletion promotes outward displacement of TM6 toward an active-like conformation. **(B)** Structural basis for monitoring receptor activation during molecular dynamics simulations. The Cα distance between R^3.50^ and R^6.30^ was used to quantify the relative positions of TM3 and TM6 as a measure of receptor activation. **(C)** Time evolution of the Cα distance between R^3.50^ and R^6.30^ in simulations of the inactive ORF74 receptor in cholesterol-free membranes and membranes containing 10%, 20%, 30%, or 40% cholesterol. The corresponding distance for the inactive ORF74 structure simulated in a cholesterol-free membrane is included for comparison. Distances were calculated by combining trajectories from all three independent simulations for each condition. **(D)** Time-dependent changes in the solvent-accessible surface area (SASA) in the simulations of the inactive ORF74 receptor in cholesterol-free membranes and membranes containing 10%, 20%, 30%, or 40% cholesterol. The corresponding SASA of the inactive ORF74 structure simulated in a cholesterol-free membrane is shown for comparison. SASA were calculated by combining trajectories from all three independent simulations for each condition. The horizontal black lines in panels C and D denote the initial TM3-TM6 distance and the SASA of the inactive ORF74 structure (t = 0 ns), respectively, and serve as reference values for evaluating time-dependent changes during the simulations.

To quantitatively evaluate these conformational changes, we monitored the Cα distance between R^3.50^ and R^6.30^ (Ballesteros-Weinstein numbering), a well-established structural measure of TM6 displacement during GPCR activation(35)(**Figure 2B**). Although ORF74 lacks the canonical ionic lock because residues 3.50 and 6.30 are both arginine, the separation between these residues remains a sensitive indicator of receptor activation(23). We next investigated whether the stabilizing effect of cholesterol was maintained over a range of physiologically relevant membrane compositions by performing additional simulations in membranes containing 10%, 20%, and 40% cholesterol. These trajectories were analyzed together with the cholesterol-free inactive and active simulations. In the absence of cholesterol, the R^3.50^-R^6.30^ distance increased progressively, approaching values characteristic of the active receptor, consistent with spontaneous TM6 opening (**Figure 2C**). In contrast, the R^3.50^-R^6.30^ distance remained remarkably stable in all cholesterol-containing membranes, indicating that cholesterol suppresses spontaneous activation over a broad range of membrane cholesterol concentrations.

Because TM6 forms one side of the cholesterol-binding pocket, its outward displacement is expected to alter the accessibility of this site. We therefore monitored the solvent-accessible surface area (SASA) of the receptor throughout the simulations. Consistent with the TM3-TM6 distance analysis, the SASA increased progressively in the cholesterol-free simulation, reflecting structural rearrangement of the pocket as the receptor transitioned toward an active-like conformation. In contrast, the pocket remained largely unchanged in membranes containing 10-40% cholesterol (**Figure 2D**), indicating that cholesterol preserves the architecture of the binding site by maintaining the inactive receptor conformation. Together, these findings demonstrate that membrane cholesterol allosterically stabilizes the inactive state of ORF74 by restricting TM6 movement and maintaining the integrity of the cholesterol-binding pocket. These findings provide a dynamic mechanistic explanation for the structural observations described above and predict that depletion of cellular cholesterol should enhance constitutive ORF74 signaling.

### The non-canonical VRY motif exposes R143^3.50^ to create an ORF74-specific cholesterol-binding pocket

To understand how ORF74 uniquely accommodates cholesterol within its transmembrane domain, we examined the structural organization of residues surrounding the cholesterol-binding pocket. A striking feature of the pocket is the direct participation of R143^3.50^, the highly conserved arginine of the class A GPCR DRY motif (**Figure 3A and B**). In most class A GPCRs, D^3.49^ forms an intramolecular ionic interaction with R^3.50^ that stabilizes the inactive receptor and regulates receptor activation(36). In ORF74, however, substitution of the conserved aspartate with valine generates a non-canonical VRY motif, eliminating this ionic interaction and fundamentally altering the structural environment surrounding R143^3.50^.

**Figure 3.**
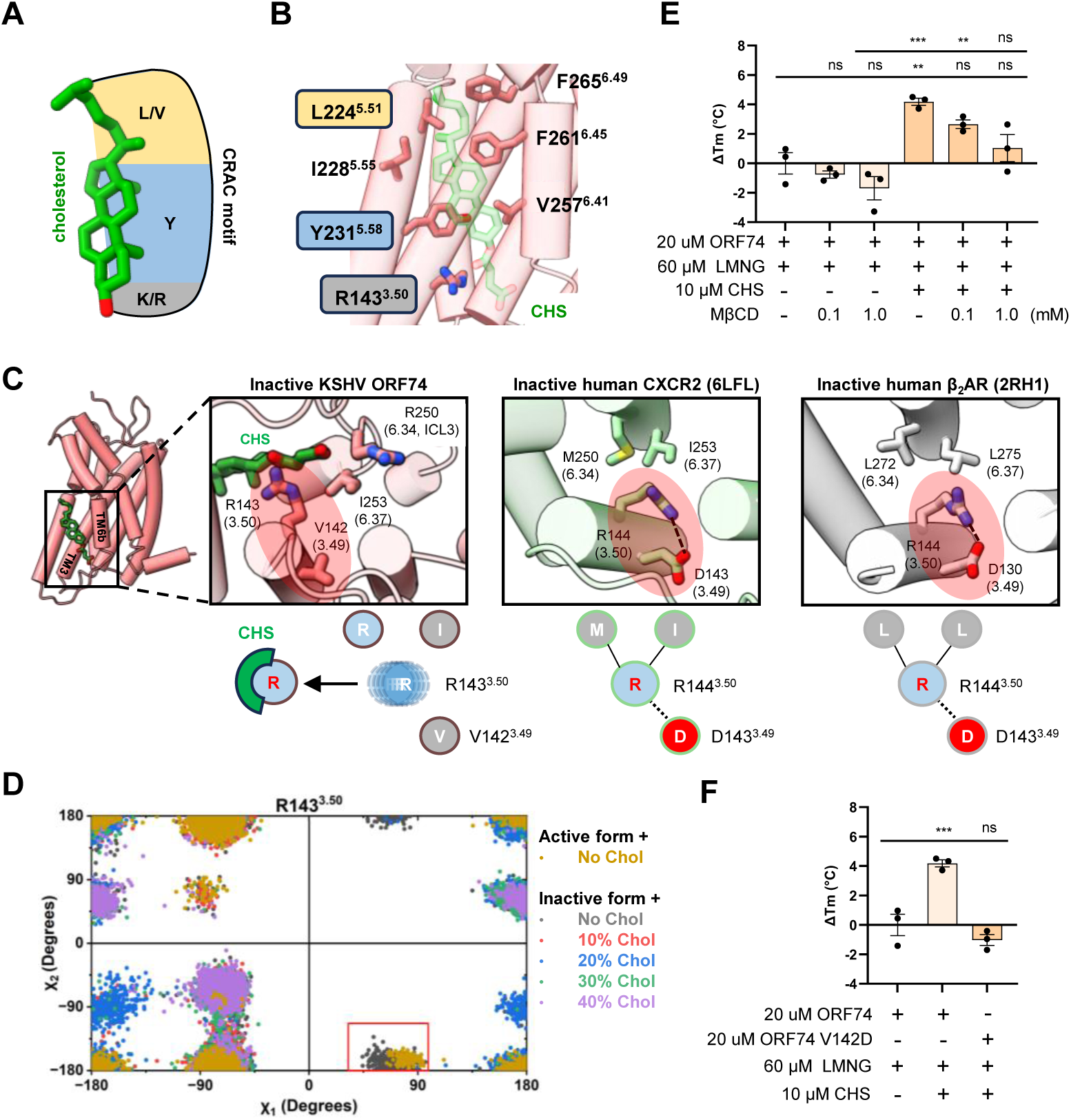
The non-canonical VRY motif enables cholesterol recognition through exposure of R^3.50^. **(A)** Schematic illustration of the canonical CRAC cholesterol-recognition motif. **(B)** Close-up view of the cholesterol-binding pocket in inactive ORF74. **(C)** Structural comparison of the DRY/VRY motifs in inactive ORF74, inactive CXCR2 (PDB: 6LFL), and inactive β2AR (PDB: 2RH1). Schematic diagrams below summarize the distinct interaction networks surrounding R^3.50^ in each receptor. **(D)** Comparison of the χ -χ dihedral angle distributions of R^3.50^ obtained from molecular dynamics simulations of the inactive and active ORF74 structures in cholesterol-free membranes and of the inactive ORF74 structure in membranes containing 10%, 20%, 30%, or 40% cholesterol. The red box highlights the rotameric states sampled by both the active receptor and the cholesterol-free inactive receptor but absent in all cholesterol-containing simulations. Rotamer distributions were generated by combining trajectories from all three independent simulations for each condition. **(E)** CHS-dependent changes in the melting temperature (ΔTm) of purified ORF74 measured by differential scanning fluorimetry (*n* = 3). ΔTm values are shown relative to ORF74 in LMNG alone. **(F)** DSF analysis comparing wild-type ORF74 and the V142D mutant (*n* = 3). Data are presented as mean ± s.e.m. from independent biological replicates. Statistical significance was determined by one-way ANOVA with multiple-comparison correction.

Structural comparison with the inactive-state structures of CXCR2 and the β2AR revealed the consequences of this substitution (**Figure 3C**). In both CXCR2 and β2AR, R^3.50^ is engaged in the canonical DRY-mediated interaction with D^3.49^, orienting its side chain toward TM3. In contrast, replacement of D^3.49^ by valine in ORF74 abolishes this interaction, leaving R143^3.50^ solvent-accessible and allowing it to adopt a distinct rotamer that projects toward the cholesterol-binding cavity (**Figure 3C, *SI Appendix* Fig. 2B)**. Consequently, R143^3.50^ directly participates in formation of the cholesterol-binding pocket together with residues contributed by TM5 and TM6, including L224^5.51^, I228^5.55^, Y231^5.58^, V257^6.34^, F261^6.38^, and F265^6.42^ (**Figure 3B**). Thus, the same evolutionary substitution that weakens the canonical inactive-state lock and promotes constitutive signaling simultaneously creates a structural interface for cholesterol recognition.

Although cholesterol-recognition motifs (CRAC/CARC) have traditionally been described as linear sequence motifs(37, 38), the ORF74 structure demonstrates that cholesterol recognition is achieved through assembly of a three-dimensional binding pocket (**Figure 3A and B**). Sequence conservation analysis revealed that R143^3.50^, Y231^5.58^, and L224^5.51^ are highly conserved among class A GPCRs, being present in approximately 94%, 75%, and 54% of receptors, respectively(39) (***SI Appendix* Fig. 2C**). Consistent with this conservation, putative CRAC-like motifs can also be identified in CXCR2 and β2AR (***SI Appendix* Fig. 2B**). However, despite the presence of sterols during structure determination, neither receptor contains cholesterol density at the corresponding position (***SI Appendix* Fig. 2D**)(32, 33). These findings indicate that conservation of CRAC-like motif-containing pocket alone is insufficient to generate a functional cholesterol-binding site and instead highlight the importance of receptor-specific three-dimensional organization. In ORF74, exposure of R143^3.50^ by the non-canonical VRY motif provides the critical structural feature that enables cholesterol recognition.

To investigate how cholesterol influences the structural organization of the cholesterol-binding pocket, we examined the rotameric behavior of R143^3.50^ during molecular dynamics simulations. In the absence of cholesterol, R143^3.50^ exhibited substantial conformational flexibility, sampling multiple χ -χ rotameric states that largely overlapped between the inactive and active receptor conformations (**Figure 3D**), consistent with our previous observations that the inactive receptor spontaneously samples active-like conformations(23). This overlap suggests that, in a cholesterol-free membrane, R143^3.50^ readily interconverts between inactive- and active-like orientations. In contrast, cholesterol binding markedly restricted the conformational landscape of R143^3.50^, confining the residue to a distinct subset of rotameric states corresponding to the inactive conformation observed in the cryoEM structure (**Figure 3D**). Notably, the rotameric states highlighted in the red box, which are populated by both the active receptor and the cholesterol-free inactive receptor, were no longer sampled in the presence of cholesterol. These findings indicate that cholesterol not only occupies the TM3-TM5-TM6 binding pocket but also allosterically limits the conformational flexibility of R143^3.50^, preventing the residue from adopting rotameric states associated with receptor activation. Together, these results identify R143^3.50^ as a cholesterol-responsive microswitch that couples cholesterol binding to stabilization of the inactive conformation of ORF74.

To experimentally validate the structural model, we first examined whether cholesterol directly stabilizes purified ORF74 using differential scanning fluorimetry (DSF). Addition of CHS increased the melting temperature (Tm) of wild-type ORF74 by approximately 4°C, indicating enhanced protein stability (**Figure 3E, *SI Appendix* Fig. 3**). This stabilizing effect was progressively reversed by methyl-β-cyclodextrin (MβCD), which sequesters cholesterol-like sterols(40), and 1 mM MβCD restored the melting temperature to that observed in the absence of CHS (**Figure 3E**). These results support a direct stabilizing interaction between cholesterol and ORF74.

We next tested whether the non-canonical VRY motif is required for cholesterol recognition by introducing the V142D substitution, thereby restoring the canonical DRY motif. We hypothesized that introduction of D142 would re-establish the intramolecular interaction with R143^3.50^, reducing its accessibility to the cholesterol-binding pocket. Consistent with this model, introduction of the V142D mutation largely eliminated the thermal stabilization normally conferred by cholesterol. CHS increased the melting temperature of wild-type ORF74 by approximately 4°C, whereas the same treatment produced little detectable stabilization of the V142D mutant (**Figure 3E and F**). Consequently, the melting temperature of the CHS-treated V142D mutant remained similar to that of untreated wild-type ORF74. These results demonstrate that restoration of the canonical DRY interaction disrupts cholesterol-dependent stabilization, supporting a model in which the non-canonical VRY motif promotes cholesterol recognition by maintaining the accessibility of R143^3.50^ within the sterol-binding pocket.

Finally, to evaluate the functional contribution of the cholesterol-binding pocket, we examined basal β-arrestin2 recruitment by ORF74 mutants carrying alanine substitutions of residues lining the cholesterol-binding cavity. Substitution of individual cholesterol-contacting residues with alanine generally reduced basal signaling relative to wild-type ORF74 **(*SI Appendix* Fig. 4)**, supporting a functional role for the cholesterol-binding pocket in maintaining normal receptor activity. In contrast, the R143A^3.50^ mutant retained wild-type basal signaling. Together, these findings suggest that the primary role of the VRY motif is to expose R143^3.50^ for cholesterol recognition, whereas the surrounding pocket residues mediate cholesterol-dependent receptor stabilization. This supports a model in which viral evolution repurposed a single structural adaptation to both promote constitutive signaling and enable host cholesterol-dependent allosteric regulation.

### Cellular cholesterol restrains constitutive ORF74 signaling

Our structural and molecular dynamics analyses predicted that membrane cholesterol stabilizes the inactive conformation of ORF74 and thereby restrains its constitutive signaling activity. To experimentally test this prediction, we reduced cellular cholesterol using BIBB-515, a selective inhibitor of oxidosqualene cyclase (OSC), a key enzyme in cholesterol biosynthesis(41) (**Figure 4A**). Because prolonged inhibition of cholesterol biosynthesis could potentially affect cell viability, we first established non-toxic concentrations of BIBB-515 for subsequent functional analyses. Treatment of HEK293T cells used for G-protein coupling assays and HTLA cells used for β-arrestin2 recruitment assays produced minimal cytotoxicity at concentrations up to 25 μM (**Figure 4B**).

**Figure 4.**
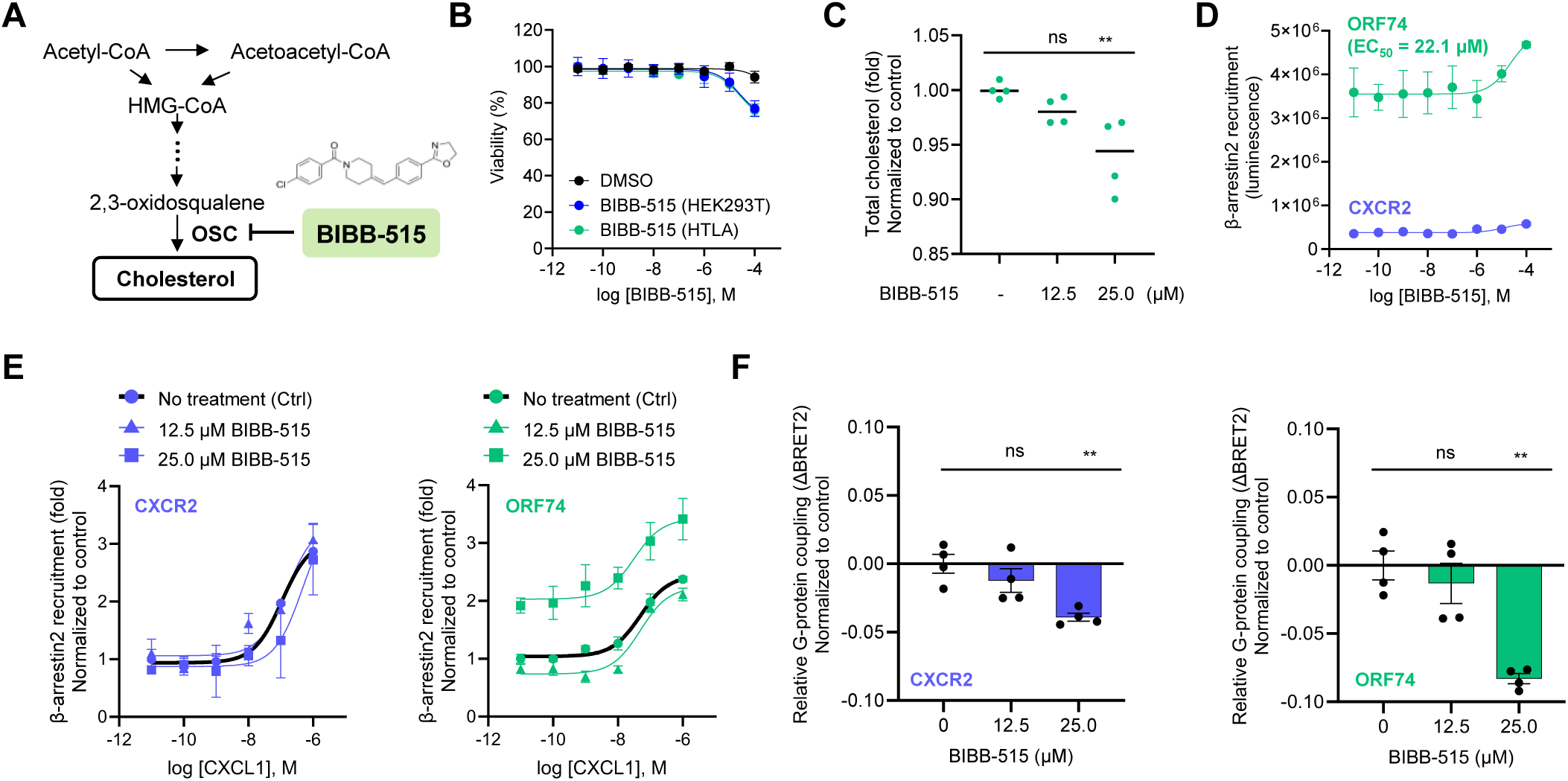
Cellular cholesterol depletion enhances constitutive ORF74 signaling. **(A)** Schematic illustration of cholesterol biosynthesis and inhibition of oxidosqualene cyclase (OSC) by BIBB-515. **(B)** Cell viability following treatment with the indicated concentrations of BIBB-515 in HEK293T and HTLA cells (*n* = 8). Cells were treated for the same duration as used in subsequent functional assays. **(C)** Relative quantification of total cellular cholesterol following treatment with BIBB-515 (*n* = 4). Cellular cholesterol levels were normalized to the untreated control. **(D)** Basal β-arrestin2 recruitment by ORF74 and CXCR2 following treatment with increasing concentrations of BIBB-515 (*n* = 3). Concentration-response analysis was used to determine the EC50 for BIBB-515-mediated enhancement of ORF74 signaling. **(E)** CXCL1-induced β-arrestin2 recruitment by CXCR2 (left) and ORF74 (right) in cells treated with the indicated concentrations of BIBB-515 (*n* = 3). Responses were normalized to the untreated control. **(F)** Relative G-protein coupling of CXCR2 (left) and ORF74 (right) following treatment with the indicated concentrations of BIBB-515 (*n* = 4). ΔBRET2 values were normalized to the untreated control. Data are presented as mean ± s.e.m. from independent biological replicates. Statistical significance was determined by one-way ANOVA with multiple-comparison correction.

We next confirmed that BIBB-515 effectively reduced cellular cholesterol levels under these experimental conditions. To exclude potential interference of the compound with cholesterol quantification, we first verified that BIBB-515 did not affect the Amplex Red cholesterol assay (***SI Appendix* Fig. 5**). Treatment with 25 μM BIBB-515 significantly reduced total cellular cholesterol compared with untreated controls (P = 0.0037), whereas 12.5 μM BIBB-515 produced only a modest reduction that did not reach statistical significance (P = 0.077) (**Figure 4C**). Thus, BIBB-515 provided two experimentally matched conditions in which cellular cholesterol was either significantly reduced or largely maintained, enabling direct assessment of the relationship between cholesterol abundance and receptor signaling.

Having established selective cholesterol depletion, we next examined whether reduced cellular cholesterol altered constitutive ORF74 signaling. β-arrestin2 recruitment was quantified using the PRESTO-Tango assay(38). BIBB-515 enhanced basal β-arrestin2 recruitment in a concentration-dependent manner, yielding an EC_50_ of approximately 22.1 μM (**Figure 4D**). To determine whether cellular cholesterol influences CXCL1-induced ORF74 activation, β-arrestin2 recruitment was measured in response to increasing concentrations of CXCL1, a well-established full agonist of ORF74(10), following treatment with BIBB-515. Treatment with 12.5 μM BIBB-515 did not significantly alter the CXCL1 concentration-response curve compared with the untreated control, indicating that this concentration had no measurable effect on agonist-induced β-arrestin2 recruitment (**Figure 4E**). In contrast, 25 μM BIBB-515 significantly increased β-arrestin2 recruitment across the entire CXCL1 concentration range, resulting in an approximately two-fold higher maximal response than the untreated control (**Figure 4E**). Notably, the concentration required to enhance receptor signaling closely matched that required to reduce cellular cholesterol, supporting a direct relationship between cholesterol depletion and ORF74 activation.

We next examined G-protein coupling using the TRUPATH BRET2 assay(42). Consistent with the β-arrestin results, treatment with 25 μM BIBB-515 significantly enhanced constitutive G-protein coupling by ORF74 (**Figure 4F**). Although a smaller increase was also observed for CXCR2, the effect was substantially greater for ORF74, consistent with a stronger modulatory effect of membrane cholesterol on ORF74 signaling. In contrast, enhanced β-arrestin2 recruitment was observed only for ORF74 and not CXCR2 (**Figure 4E**), further indicating that cholesterol-dependent regulation of β-arrestin signaling is a distinctive feature of ORF74. Importantly, the cellular responses closely paralleled the extent of cholesterol depletion. Treatment with 12.5 μM BIBB-515 neither significantly reduced cellular cholesterol nor altered receptor signaling, whereas 25 μM BIBB-515 both lowered cellular cholesterol and significantly enhanced β-arrestin2 recruitment and G-protein coupling (**Figure 4E and F**). The close correspondence between cholesterol depletion and receptor activation argues against nonspecific drug effects and instead supports a direct functional role for membrane cholesterol in restraining constitutive ORF74 signaling.

Collectively, these cellular experiments validate the structural, molecular dynamics, and biochemical findings presented above. Together, our data support a unified model in which host membrane cholesterol functions as an endogenous allosteric regulator that restrains constitutive ORF74 signaling by stabilizing the inactive receptor conformation.

## DISCUSSION

In this study, we identify a previously unrecognized cholesterol-dependent allosteric mechanism that restrains constitutive signaling of the KSHV-encoded GPCR ORF74. By integrating cryoEM structural analysis, molecular dynamics simulations, biochemical characterization, and cellular signaling assays, we show that membrane cholesterol binds a unique TM3-TM5-TM6 pocket present only in the inactive receptor conformation (**Figure 1**). Cholesterol binding restricts the conformational flexibility of TM6, stabilizes the inactive state, and suppresses spontaneous receptor activation (**Figure 2**). Molecular dynamics simulations further revealed that, in the absence of cholesterol, the inactive receptor spontaneously undergoes an outward displacement of TM6 toward an active-like conformation, accompanied by remodeling of the cholesterol-binding pocket. In contrast, this transition was consistently suppressed in membranes containing physiological levels of cholesterol (10-40%), indicating that membrane cholesterol reshapes the conformational landscape of ORF74 by biasing the receptor toward its inactive state rather than simply stabilizing a single static structure. These findings resolve the longstanding paradox arising from our previous structural studies, in which ORF74 exhibited exceptionally high constitutive signaling despite adopting a canonical inactive conformation in the absence of ligand(23).

A central finding of this work is that cholesterol recognition is enabled by the non-canonical VRY motif that distinguishes ORF74 from most class A GPCRs. In canonical receptors, the conserved D^3.49^ residue forms an ionic interaction with R^3.50^, restricting the orientation of this highly conserved arginine within the DRY motif. Replacement of D^3.49^ with valine abolishes this interaction, exposing R143^3.50^ toward the membrane-facing cavity where it directly participates in cholesterol recognition (**Figure 3**). Rather than functioning through a conventional linear CRAC sequence, ORF74 uniquely assembles a three-dimensional cholesterol-binding pocket in which R143^3.50^, together with residues from TM5 and TM6, forms a receptor-specific structural interface (**Figure 3B**). Thus, the VRY motif not only distinguishes ORF74 from cellular GPCRs but also couples membrane cholesterol to receptor conformational control through a previously unrecognized allosteric mechanism.

Our findings also extend current understanding of cholesterol-mediated GPCR regulation. Cholesterol has been shown to influence numerous cellular GPCRs by modulating membrane organization, ligand binding, receptor oligomerization, and conformational stability(29). In most cases, however, cholesterol has been viewed as a permissive membrane component or a modulator of ligand-dependent receptor activation. In contrast, ORF74 utilizes cholesterol as an endogenous allosteric regulator that restrains constitutive receptor activity. The location of the cholesterol-binding pocket immediately adjacent to TM6, the principal conformational switch of class A GPCR activation, places cholesterol in a unique position to directly control spontaneous receptor activation. Although additional studies will be required to determine whether analogous mechanisms exist in other viral or cellular GPCRs, our findings establish that membrane cholesterol can directly regulate constitutive receptor signaling through stabilization of the inactive receptor conformation.

The biological implications of this mechanism may extend beyond ORF74. Persistent viruses are unlikely to benefit from simply maximizing signaling output. Although constitutive signaling promotes viral persistence, proliferation, inflammation, angiogenesis, and oncogenic transformation, unrestricted receptor activation would also be expected to compromise viral fitness by disrupting cellular homeostasis, increasing immune surveillance, or reducing the long-term survival of infected cells. Our findings suggest that KSHV addresses this challenge by coupling constitutive ORF74 signaling to a host-derived membrane lipid. Rather than acting solely as a structural membrane component, cholesterol functions as an endogenous allosteric regulator that constrains excessive receptor activation while preserving constitutive signaling. This mechanism provides a plausible strategy by which persistent viruses can maintain sustained signaling while avoiding detrimental overactivation.

An intriguing aspect of this mechanism is that the structural adaptation previously associated with constitutive signaling also creates a cholesterol-responsive allosteric interface. Our previous work established that replacement of the highly conserved DRY motif with a non-canonical VRY motif contributes to the exceptional constitutive activity of ORF74(23). Here, we show that the same structural rearrangement exposes R143^3.50^ to generate a cholesterol-binding interface that links receptor activity to the host membrane environment. Molecular dynamics simulations further revealed that cholesterol not only occupies this pocket but also restricts the rotameric flexibility of R143^3.50^, preventing the residue from sampling conformations associated with receptor activation. These observations identify R143^3.50^ as a cholesterol-responsive microswitch that dynamically couples sterol binding to stabilization of the inactive receptor conformation. Rather than functioning as opposing regulatory mechanisms, these two properties appear to be structurally coupled: the VRY motif increases the intrinsic signaling propensity of ORF74, while simultaneously rendering that activity responsive to membrane cholesterol. This coupling provides a simple structural solution by which constitutive signaling can remain dynamically regulated by the host membrane. The exact mechanism underlying this coupling remains to be determined and is likely to be complex. For example, our simulation study of a different system, the EphA2 receptor dimer showed that although the membrane is generally thickened by cholesterol, thus disfavoring displacement of helices away from a membrane normal, actually a local membrane thinning was observed in those simulations around the protein in cholesterol(43).

More broadly, our findings support the concept that persistent viruses may not evolve to maximize signaling activity, but rather to optimize it. By exploiting host membrane cholesterol as an allosteric regulator, ORF74 maintains constitutive signaling while preserving the capacity for structural restraint through the inactive receptor state. Whether similar host lipid-dependent regulatory mechanisms have evolved in other viral GPCRs remains an important question for future investigation. Beyond viral receptors, these findings suggest that membrane cholesterol may represent a broader mechanism for regulating constitutively active GPCRs and identify the ORF74 cholesterol-binding pocket as a potential therapeutic target for KSHV-associated malignancies.

## MATERIALS AND METHODS

### Structure refinement

The cryoEM map EMD-43717 (https://www.ebi.ac.uk/emdb/EMD-43717) was used for model building of KSHV ORF74 and cholesterol. The atomic coordinates have been deposited in the Protein Data Bank (PDB) under accession codes 36XZ (https://www.rcsb.org/structure/36XZ). The Q-scores for model building are provided in ***SI Appendix* Fig. 1A**. Model building was performed in Coot(44), and figures were prepared using PyMOL and ChimeraX(45).

### Differential scanning fluorimetry assay

To determine the melting temperature of the protein, LMNG-solubilized purified ORF74 was mixed with BODIPY FL L-cystine (ThermoFisher, B20340), and temperature-dependent fluorescence changes were measured using a qPCR instrument (Biorad, C1000 Touch Thermal Cycler). The protein and dye concentrations, total sample volumes, and instrument settings were based on previously published protocols(46). The resulting fluorescence data were analyzed using the Boltzmann (sigmoidal) nonlinear regression model implemented in GraphPad Prism (version 10.1.2).

### Total cholesterol measurement

Intracellular total cholesterol levels were measured using a commercially available Amplex™ Red Cholesterol Assay Kit (Invitrogen, A12216) according to the manufacturer’s instructions. HEK293T cells (ATCC, CRL-3216) cultured in DMEM (Gibco, 11965092) supplemented with 10% (v/v) fetal bovine serum (FBS, Gibco, A5256701) and 1% (v/v) penicillin-streptomycin (10,000 U/mL, Gibco, 15140122) (complete medium) were used for all experiments. Cells were transfected with CXCR2 or ORF74 expression constructs using the TransIT-2020 reagent (Mirus Bio, MIR 5404) in tissue culture-treated 6-well plates (Corning, 353046). After 16-20 h, the medium was replaced with a fresh complete medium and cells were incubated for an additional 24 h. Cells were then treated with BIBB-515 (Cayman, 10010517) and harvested 16 h later. Harvested cells were lysed in PBS containing 1% Triton X-100 and heated at 95°C for 5 min. After centrifugation at 13,000 rpm for 10 min, the supernatants were collected and subjected to the Amplex™ Red cholesterol assay.

### Cell viability measurement

Cell viability was assessed using a commercially available CellTiter-Glo® 2.0 Cell Viability Assay (Promega, G9241) according to the manufacturer’s instructions. HTLA (a HEK293 cell line stably expressing a tTA-dependent luciferase reporter and a β-arrestin2-TEV fusion gene) and HEK293T cells cultured in complete DMEM were seeded into cell culture-treated 96-well plates (Corning, 3917) and incubated overnight. The medium was then removed, and cells were washed twice with FBS-free FreeStyle 293 expression medium (Gibco, 12338018) followed by replacement with the same medium. Cells were treated with BIBB-515 or 25-hydroxycholesterol (MedChemExpress, HY-113134) in a dose-dependent manner and incubated for 16 h. Cell viability was subsequently measured using the CellTiter-Glo® 2.0 assay.

### β-Arrestin2 recruitment assay (PRESTO-Tango assay)

HEK293-derived HTLA cells stably harboring a tTA-driven luciferase reporter and a β-arrestin2-TEV fusion construct were generously provided by the laboratory of Brian Roth. Cells were cultured in DMEM supplemented with 10% FBS, 100 U/mL penicillin, 100 μg/mL streptomycin. GPCRs including ORF74 and CXCR2 constructs were generated using the Tango plasmid system (Addgene kit 1000000068) and introduced into the corresponding expression vectors. Transfections were performed in 6-well plates using the TransIT-2020 reagent (Mirus Bio, MIR5400) and allowed to proceed overnight. Cells were subsequently detached with a trypsin-free dissociation reagent and seeded onto poly-L-lysine-coated white 96-well plates (ThermoFisher Scientific, 15042) at a density of 8,000-10,000 cells per well in FreeStyle 293 expression medium (Gibco, 12338018). After 24 h, drug treatment was carried out using a concentration gradient, followed by an additional 18 h incubation. Luminescence was then quantified by adding Bright-Glo reagent (Promega, E2610) to each well and incubating for 15 min at room temperature. Signal intensity was measured using a Varioskan LUX plate reader (ThermoFisher Scientific). Data analysis was performed with GraphPad Prism (version 10.1.2).

### G protein coupling assay (TRUPATH assay)

Rluc8-tagged Gα and GFP2-tagged Gγ expression constructs were acquired from Addgene (Kit #1000000163) and utilized according to the TRUPATH protocol(42). HEK293T cells (ATCC, CRL-3216) were seeded in 6-well culture plates (Corning, 3516) and transiently transfected using TransIT-2020 reagent (Mirus Bio, MIR5400) in DMEM supplemented with 10% FBS, 100 U/mL penicillin, 100 μg/mL streptomycin. The transfection mixture comprised plasmids encoding the ORF74 or CXCR2 together with Rluc8-fused Gα, Gβ, and GFP2-fused Gγ subunits. At 24 h post-transfection, cells were harvested and replated into white, opaque 96-well plates (ThermoFisher Scientific, 15042), followed by replacement of the culture medium with FreeStyle 293 expression medium (Gibco, 12338018). After an additional 24 h incubation, BRET signal measurements were conducted. Recombinant CXCL1 (Biotechne, 275-GR) was applied as the ligand. Coelenterazine 400a (Goldbio, C-320-1), prepared in ethanol, was added to each well at a final concentration of 50 μM with thorough mixing to ensure homogeneity. BRET signals were acquired using an EnVision 2105 plate reader equipped with a BRET2 dual-emission optical module (Revvity, 2100-4150). Data processing and analysis were performed using GraphPad Prism (version 10.1.2).

### Surface expression analysis by FACS

Cells expressing N-terminally FLAG-tagged GPCRs were harvested and washed with FACS buffer (DPBS containing 2% BSA). Cell-surface receptors were stained with APC-conjugated anti-FLAG antibody (Abcam, ab72569) for 30 min at 4 °C. Following three washes with FACS buffer, cells were resuspended in the same buffer and analyzed using a BD FACSCelesta flow cytometer (BD Biosciences). Flow cytometry data were processed and quantified using FlowJo software (version 10; BD Biosciences).

### Statistical analysis

Data are presented as mean ± SEM. Statistical analyses were performed using GraphPad Prism (version 10.1.2). Comparisons among multiple groups were analyzed by one-way ANOVA followed by Dunnett’s multiple comparisons test. Depending on the experiment, ORF74 WT or CXCR2 WT served as the reference group for statistical comparisons. Differences were considered statistically significant at *P* < 0.05 (\*\*\*\**P* < 0.0001, \*\**P* < 0.001, \**P* < 0.01, *P* < 0.05; ns, *P* ≥ 0.05). Sample sizes ranged from *n* = 2 to *n* = 8 depending on the assay, and the exact *n* values are provided in the corresponding figure legends.

### System setup for molecular dynamics simulations

The inactive ORF74 cryoEM structure(23) was used as the starting model for all simulations. To investigate the effect of membrane cholesterol on receptor dynamics, ORF74 was embedded in POPC bilayers containing 10%, 20%, 30%, or 40% cholesterol using the CHARMM-GUI Membrane Builder(47). For comparison, simulation data from our previous study, including the inactive ORF74 receptor in a cholesterol-free membrane and the active ORF74 structure bound to CXCL1 and the C-terminal α5 helix (K330-F354) of the Gαi subunit, were included for analysis(23). Each system was solvated with TIP3P water, and Na^+^ and Cl^-^ions were added to neutralize the system and achieve a final salt concentration of 150 mM. The resulting systems contained approximately 73,000-82,000 atoms. All simulations were performed using GROMACS 2023.3 with the CHARMM36m force field. Long-range electrostatic interactions were calculated using the particle mesh Ewald (PME) method, and a 1.2 nm cutoff was applied for short-range nonbonded interactions. Covalent bonds involving hydrogen atoms were constrained using the SHAKE algorithm. Temperature and pressure were maintained using the Nosé-Hoover thermostat and Parrinello-Rahman barostat, respectively. Each system was energy-minimized using the steepest descent algorithm, followed by six sequential equilibration steps with gradually released positional restraints prior to production simulations.

### Well-tempered metadynamics simulations

Enhanced sampling simulations were performed using PLUMED 2.9.0 interfaced with GROMACS 2023.3(48). Two collective variables (CVs) were employed to monitor receptor activation: (i) the distance between the centers of mass of the TM3b and TM6b regions and (ii) the backbone root-mean-square deviation (RMSD) of the receptor relative to the inactive structure. Well-tempered metadynamics bias potentials were deposited every 500 integration steps with an initial hill height of 0.2 kcal/mol and a bias factor of 5. Simulations were continued until sufficient sampling of the conformational landscape was achieved. Each system was simulated for 200 ns, and three independent simulations were performed for every cholesterol condition, yielding an aggregate sampling time of 600 ns per system.

### Trajectory analysis

Simulation trajectories were analyzed using the built-in GROMACS analysis tools(48). Structural stability and receptor activation were evaluated by monitoring the Cα distance between residues R^3.50^ and R^6.30^, solvent-accessible surface area (SASA) of the cholesterol-binding pocket, and the χ1 and χ2 dihedral angle distributions. Structural snapshots were generated using PyMOL, and all quantitative analyses were plotted using Origin 2022.

## DATA, MATERIALS, and SOFTWARE AVAILABILITY

The atomic coordinates for cholesterol bound ORF74 have been deposited in the RCSB PDB 36XZ (https://www.rcsb.org/structure/36XZ).

## ACKNOWLEDGEMENTS

This work was supported by the National Institutes of Health through grants CA251275 (J.J.) and AG084065, EY029169, and AG089561 (M.B.). The work of ARS and MB utilized the high-performance computing resources at CWRU.

## AUTHOR INFORMATION

These authors contributed equally to this work: Jun Bae Park, Amita Rani Sahoo.

### Contributions

J.B.P. and J.U.J. conceived the study, designed the research, and participated in all aspects of the project, including hypothesis development and experimental validation. J.B.P. performed structural, biochemical, and cell-based functional analyses. A.R.S. and M.B. planned, executed and analyzed all the MD simulation-related data. A.R. contributed to experimental design, scientific discussions, and experimental work. J.B.P., A.R.S., and J.U.J. wrote the manuscript. All authors reviewed the final manuscript.

### Corresponding author

Correspondence to Jun Bae Park and Jae U. Jung.

## ETHICS DECLARATIONS

### Competing interests

The authors declare no competing interests.

---

**SI Appendix Fig. 1.**
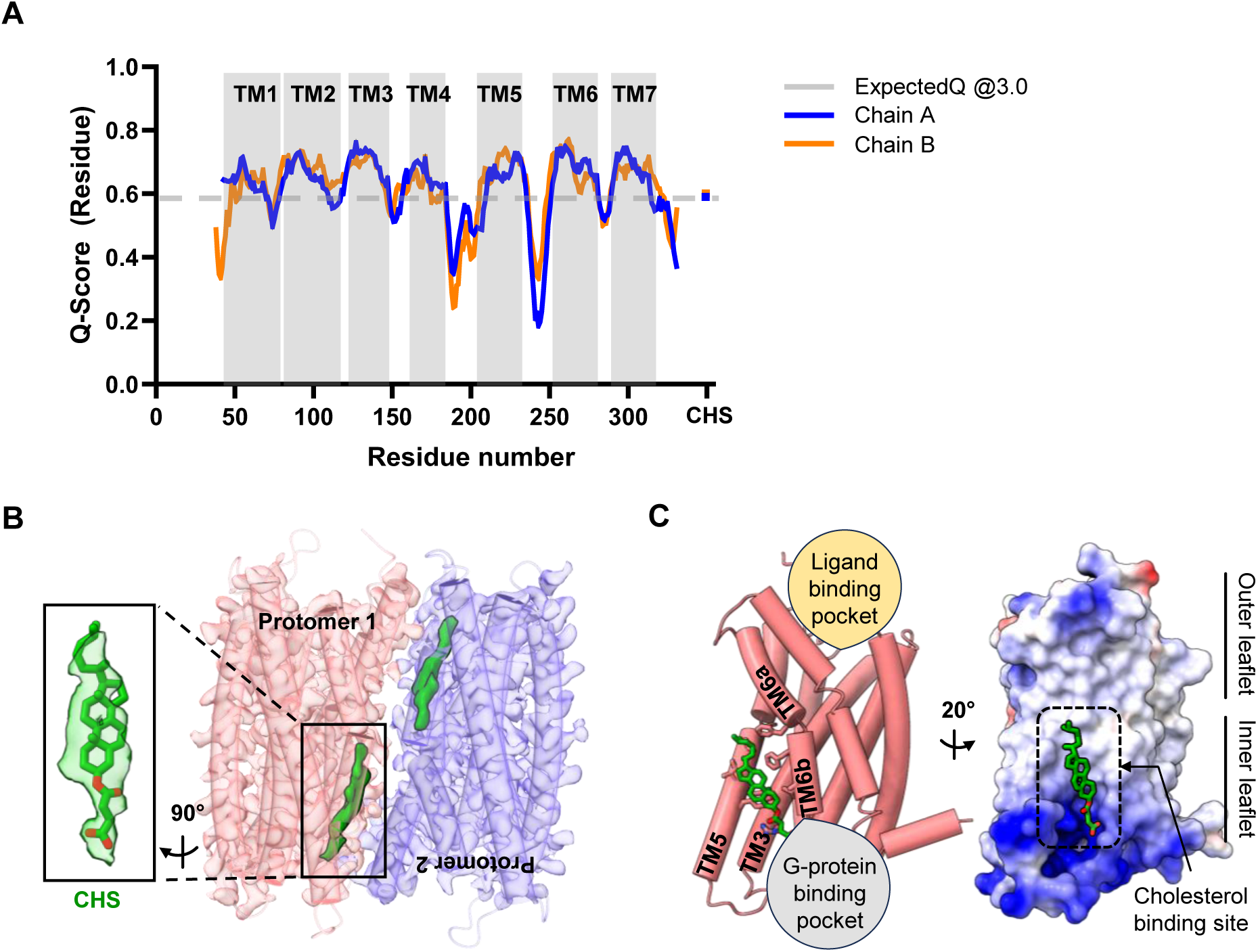
**(A)** Per-residue Q-score profile of the inactive ORF74 cryoEM structure. Q-scores are shown for Chain A (blue) and Chain B (orange). Gray shaded regions indicate TM helices (TM1-TM7), and the dashed line denotes the expected Q-score at 3.0 Å resolution. The Q-score of the modeled CHS is shown on the right. **(B)** CryoEM structure of the inactive ORF74 dimer showing an additional density assigned as cholesteryl hemisuccinate (green) at each protomer. **(C)** Location of the CHS-binding pocket within ORF74. Left, side view of the inactive ORF74 structure highlighting the CHS molecule (green) positioned between TM3, TM5, and TM6. Right, electrostatic surface representation showing that the cholesterol-binding site is located on the inner membrane leaflet, spatially separated from the extracellular ligand-binding pocket.

**SI Appendix Fig. 2.**
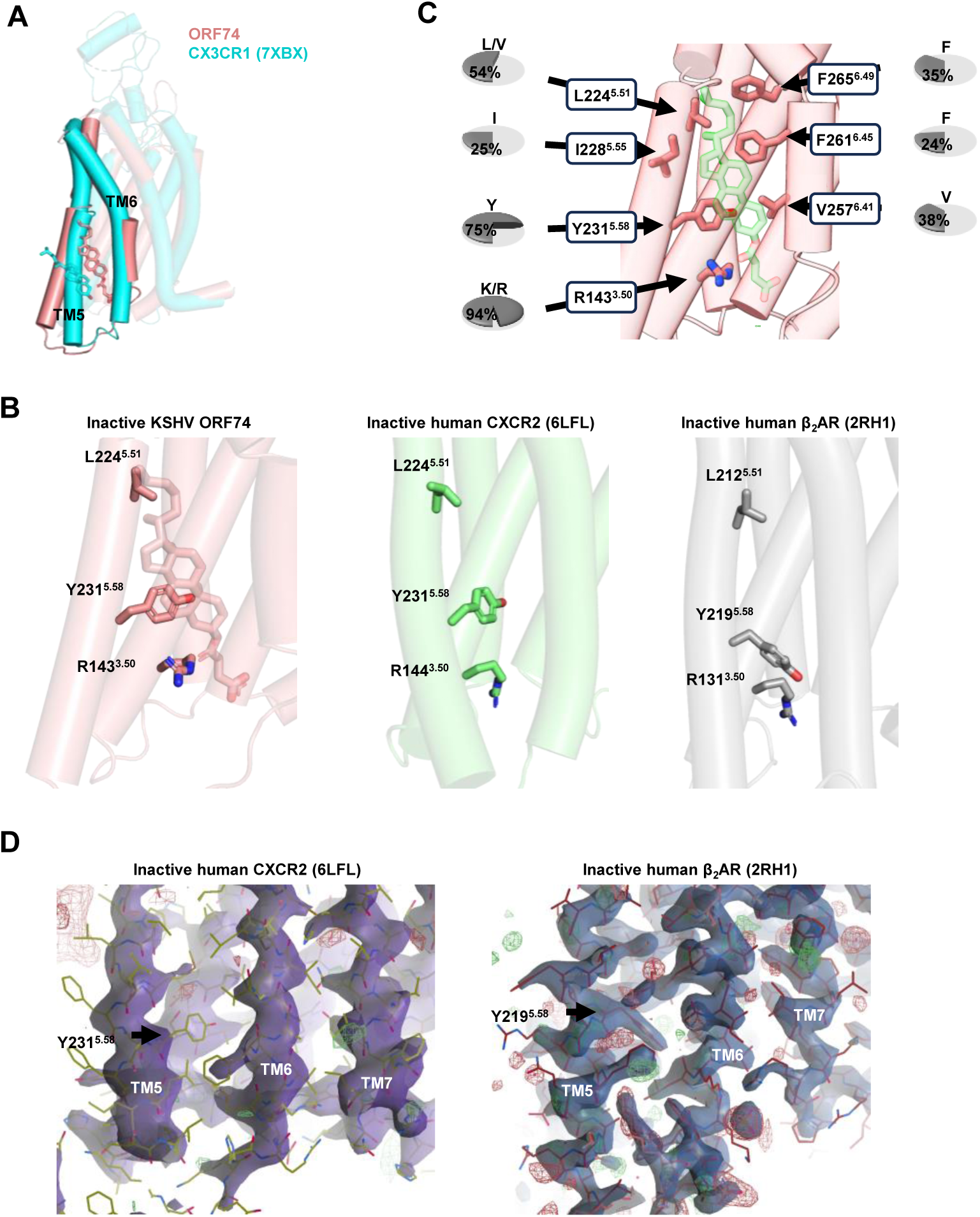
**(A)** Structural superposition of inactive ORF74 (salmon) and inactive CX3CR1 (cyan, PDB: 7XBX). The cholesterol-binding pocket in ORF74 is formed by TM3, TM5, and TM6, whereas the cholesterol-binding site in CX3CR1 is located at a distinct position. **(B)** Comparison of the R3.50 rotamer conformation in inactive ORF74, inactive CXCR2 (PDB: 6LFL), and inactive β2AR (PDB: 2RH1). **(C)** Close-up view of the cholesterol-binding pocket in inactive ORF74. Sequence conservation of the corresponding residues among ORF74 homologs is indicated. **(D)** Structural comparison of the region corresponding to the ORF74 cholesterol-binding pocket in inactive CXCR2 (PDB: 6LFL) and inactive β2AR (PDB: 2RH1).

**SI Appendix Fig. 3.**
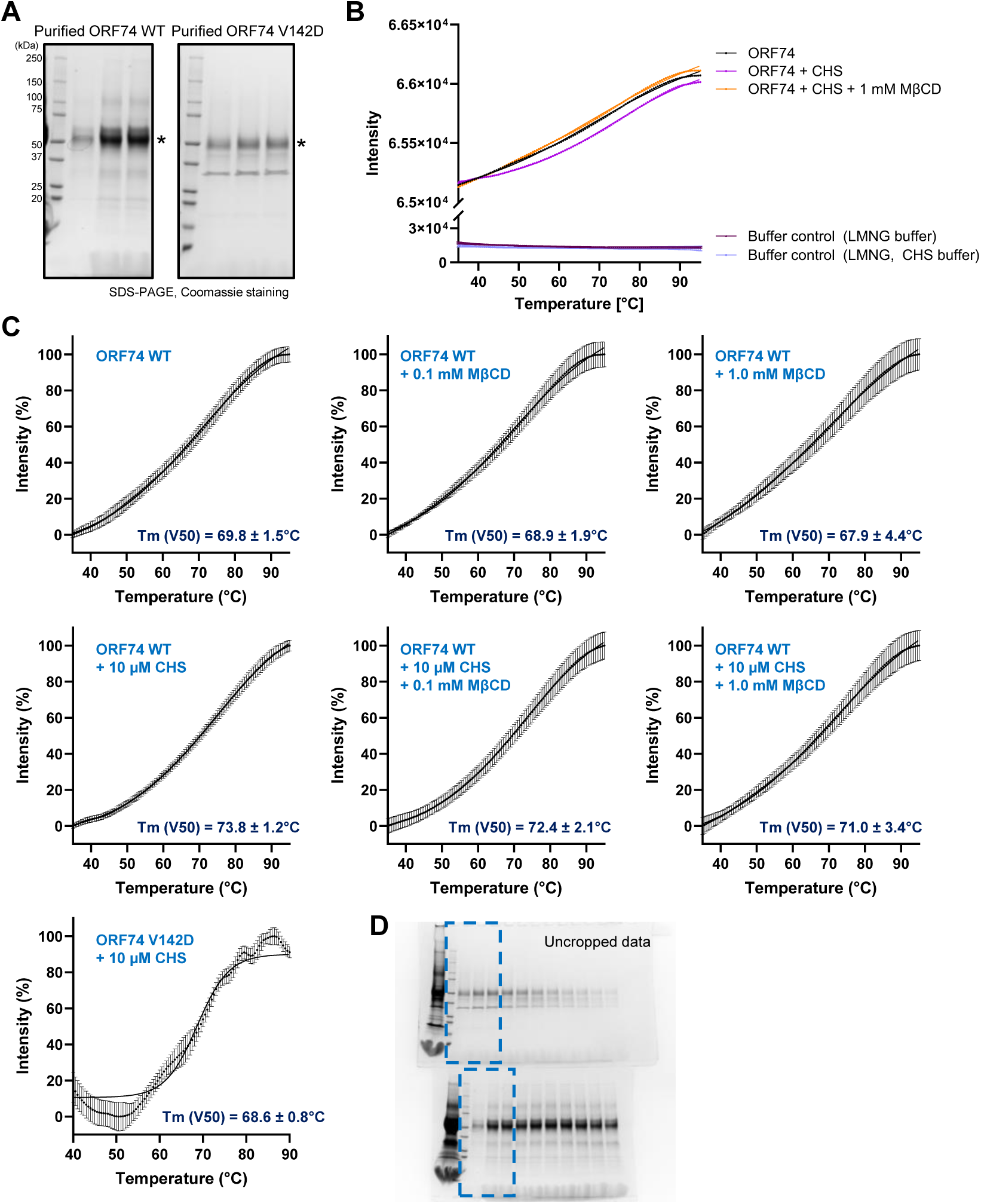
Differential scanning fluorimetry (DSF) analysis of purified ORF74. **(A)** Purified wild-type ORF74 and the V142D mutant analyzed by Coomassie-stained SDS-PAGE. Asterisks indicate the ORF74 protein bands. **(B)** Preliminary DSF fluorescence profiles used to assess the effects of CHS and MβCD on purified ORF74. Representative curves for ORF74 in LMNG, LMNG containing 10 μM CHS, and LMNG containing 10 μM CHS plus 1.0 mM MβCD are shown together with the corresponding buffer controls. **(C)** Representative DSF melting curves under the indicated conditions (*n* = 3). Melting temperatures (Tm, V50) were determined for wild-type ORF74 in LMNG alone, LMNG containing 10 μM CHS, and LMNG containing CHS with 0.1 or 1.0 mM MβCD. The V142D mutant was analyzed in LMNG containing 10 μM CHS. Melting curves were fitted using the built-in Boltzmann sigmoidal model in GraphPad Prism, and Tm (V50) values were calculated from the fitted curves. **(D)** Representative uncropped SDS-PAGE gel corresponding to the purified proteins used for DSF experiments. Dashed boxes indicate the regions shown in panel (A).

**SI Appendix Fig. 4.**
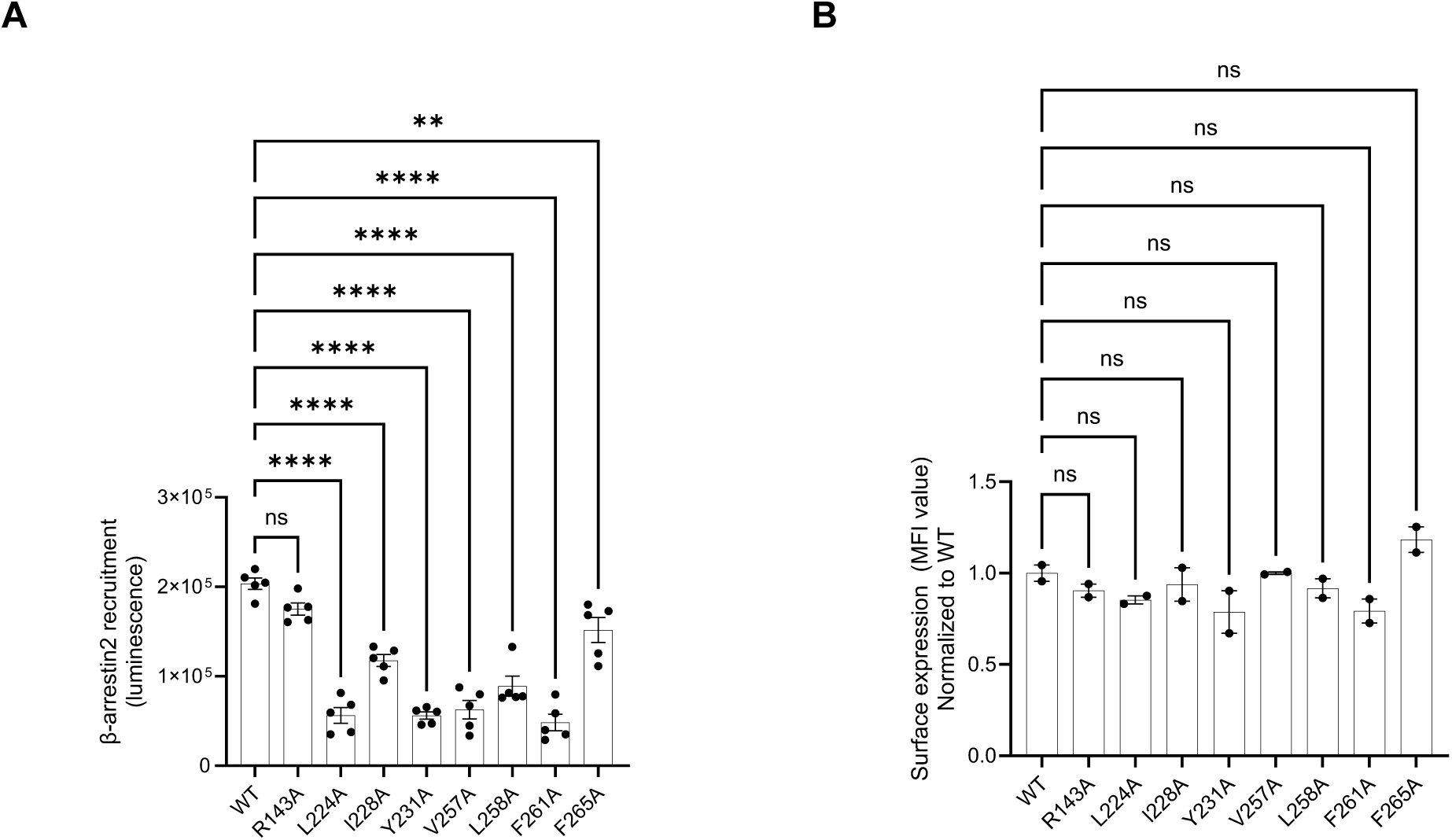
Functional validation of residues comprising the cholesterol-binding pocket. **(A)** Basal β-arrestin2 recruitment by wild-type ORF74 and cholesterol-binding pocket mutants measured using the PRESTO-Tango assay (*n* = 5). Luminescence signals represent constitutive receptor activity. Data are presented as mean ± s.e.m. from independent biological replicates. Statistical significance was determined by one-way ANOVA with multiple-comparison correction. **(B)** Cell-surface expression of wild-type ORF74 and the indicated mutants measured by flow cytometry (*n* = 2). Mean fluorescence intensity (MFI) was normalized to wild-type ORF74. Data are presented as mean ± s.e.m. from independent biological replicates. Statistical analysis was performed by one-way ANOVA with multiple-comparison correction.

**SI Appendix Fig. 5.**
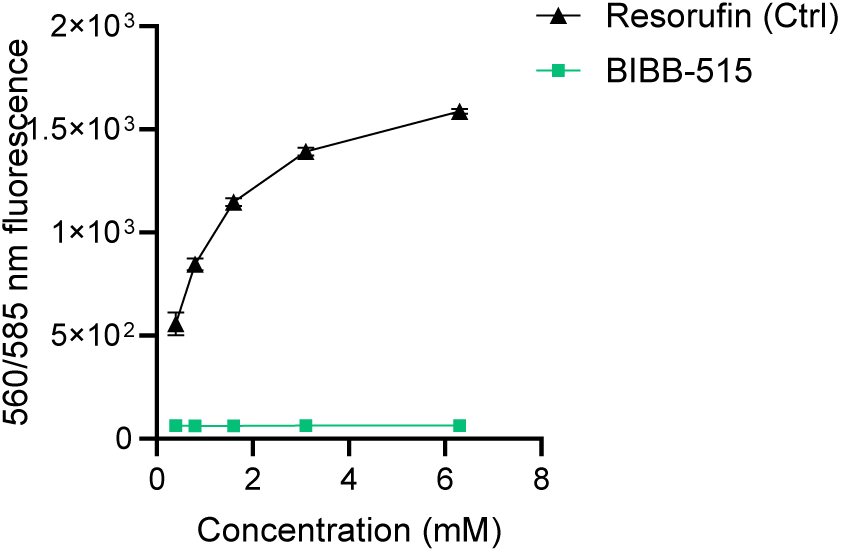
BIBB-515 does not interfere with the Amplex Red cholesterol assay. Increasing concentrations of resorufin (positive control), BIBB-515 was incubated with the Amplex Red cholesterol detection reagents in the absence of cells (*n* = 3). Fluorescence (560/585 nm) was measured to evaluate potential assay interference. Resorufin produced a concentration-dependent increase in fluorescence, whereas BIBB-515 showed no detectable fluorescence under the same conditions.

## Notes

### Competing Interest Statement

The authors have declared no competing interest.

